# Role of TM3 in claudin-15 strand flexibility: a molecular dynamics study

**DOI:** 10.1101/2022.06.08.494751

**Authors:** Shadi Fuladi, Sarah McGuinness, Fatemeh Khalili-Araghi

## Abstract

Claudins are cell-cell adhesion proteins within tight junctions that connect epithelial cells together. Claudins polymerize into a network of strand-like structures within the membrane of adjoining cells and create ion channels that control paracellular permeability to water and small molecules. Tight junction morphology and barrier function is tissue specific and regulated by claudin subtypes. Here, we present a molecular dynamics study of claudin-15 strands within lipid membranes and the role of a single-point mutation (A134P) on the third transmembrane helix (TM3) of claudin-15 in determining the morphology of the strand. Our results indicate that the A134P mutation significantly affects the lateral flexibility of the strands, increasing the persistence length of claudin-15 strands by a factor of three. Analyses of claudin-claudin contact in our *µ*second-long trajectories show that the mutation does not alter the intermolecular contacts (interfaces) between claudins. However, the dynamics and frequency of interfacial contacts are significantly affected. The A134P mutation introduces a kink in TM3 of claudin-15 similar to the one observed in claudin-3 crystal structure. The kink on TM3 skews the rotational flexibility of the claudins in the strands and limits their fluctuation in one direction. This asymmetric movement in the context of the double rows reduces the lateral flexibility of the strand and leads to higher persistence lengths of the mutant.

## Introduction

Tight junctions (TJs) are macromolecular structures that connect the apical surface of epithelial cells. In freeze-fracture electron microscopy, they appear as a network of linear strands in the cell membrane^1–4^, and form a barrier to control transport of small molecules across epithelia^5–8^. Disruption of TJ and their barrier function can lead to numerous diseases pertaining to liquid retention in tissues^9,10^. As key components of tight junctions, claudins polymerize into strands which regulate paracellular permeability across epithelial tissue layers^8,11–15^. So far, 27 members (subtypes) of the claudin family are identified in mammals^12^. Claudin subtypes express subtype-specific functional differences as well as subtype-specific strand morphology, and can even assemble into heterogenous strands with other subtypes^13,16,17^. Although the mechanism of assembly is still a mystery, our understanding of claudin strand assembly has improved by the structural insights revealed by recent crystallographic structures^18–21^, and the functional models derived for claudin channels^15,22–26^.

Claudins are tetra-span membrane proteins with two extracellular segments, ECS1 and ECS2, which form a five-strand *β*-sheet close to the surface of the membrane^18,20,21,24,27^. Claudin strands in TJs are formed by *cis-*interactions between monomers in the same membrane and *trans-*interactions between extracellular segments of monomers between two cells^28–32^. One of the models for claudin strand polymerization, inspired by the crystallographic structure of mouse claudin-15 and cysteine cross-linking experiments, proposed that claudins assemble into an anti-parallel double row within the membrane which is then attached to a similar structure on an adjacent cell^24^. This architecture of claudin strands originally proposed by Suzuki et al^24^ was later verified by molecular dynamics (MD) simulations and docking studies of claudin-15 to generate plausible architectures of claudin strands, including evidence for cation-selective claudin pore transport^22,23,33–35^.

In this model, the *cis-*interactions involve side-by-side interactions of claudins at the extracellular side involving the third transmembrane helix TM3, ECS2 and a short extracellular helix ECH that is parallel to the membrane^22–24,28–30^. Critical to the proposed architecture is another set of *cis-*interactions between two anti-parallel rows of claudins in the membrane leading to “face-to-face” dimerization of *β*-sheets to generate the pipe-like structure of claudin pores^24^. Studies of claudin strands in fibroblasts show that claudin strands are dynamic with the ability to arch, bend, and form new branches through remodeling^35–37^. However, the molecular basis for claudin strand branching, bending, and remodeling remains unaccounted for in this model^24^.

We recently developed a mechanistic model that describes the molecular nature of strand flexibility within this context^38^. In our model, claudins form interlocking tetrameric ion channels that slide with respect to each other to accommodate local curvature of the strand without loss of function. Simulations of claudin-15 strands at large scale showed that this movement is facilitated by flexible side-by-side *cis-*interactions that allow rotation and displacement of claudin monomers with respect to each within the membrane^38^.

Recent crystallographic structures of claudin-3 and two of its mutants P134A and P134G suggests that *cis-*interactions between claudins is indirectly affected by a single mutation at position 134 on the third transmembrane helix (TM3) of claudin-3^21^. In members of the mammalian claudin family, this position is mainly occupied by glycine, proline (as in claudin-3), or alanine (as in claudin-15). With a protein structure very similar to claudin-15, TJ strands formed by claudin-3 showed a distinct morphology with sparsely distributed almost linear strands. On the other hand, the TJ strands formed by two claudin-3 mutants, P134A and P134G, appeared to be highly flexible with many hairpin curves similar to those formed by claudin-15. Structural comparison of wild-type (WT) claudin-3 with its mutants indicates that the proline residue at position 134 bends the third transmembrane helix (TM3) toward the membrane pulling the entire extracellular domain toward the membrane by approximately 8^°^. Intriguingly, mutations of P134 to a glycine or alanine recovered the highly flexible TJ morphology with many hairpin curves, similar to the TJs formed by claudin-15. It was therefore suggested that the kink in the secondary structure of TM3 might limit the mobility of claudins in the strand by indirectly stabilizing the *cis-*interactions between them^21^. However, this remains to be seen in claudin-15.

Here, we present a molecular dynamics (MD) study of claudin-15 strands in lipid membranes and compare mechanical properties of wild-type (WT) and A134P mutant strands. We show that the A134P mutation significantly affects the lateral flexibility and the persistence length of the strands. Moreover, our analyses of the simulation trajectories and side-by-side *cis-*interactions between claudins reveal the molecular mechanism for the increased stiffness of the mutant strands. We used a combination of all-atom and hybrid resolution models for our simulations. The allatom simulations were used as benchmarks and control simulations to establish the stability of the models and to identify inter-atomic interactions with more accuracy. On the other hand, the hybrid resolution models^38^, while preserving the atomic nature of the protein, allowed us to study the dynamics of the strands at large scales (sub-*µ*m) and at long time scales required for persistence length calculations.

## Materials and methods

### System setup

Four sets of claudin-15 strand models were prepared for molecular dynamics (MD) simulations. Two systems of “single layer” claudin-15 strands corresponding to wild-type and the A134P mutant in a POPC lipid bilayer were constructed and simulated with atomic resolution. In addition, two sets of claudin-15 strands in a double membrane system were constructed for the wild-type (WT) and A134P mutant at various lengths (30 nm to 135 nm) and were simulated at hybrid resolution as described below (Fig. 1).

**Figure 1:**
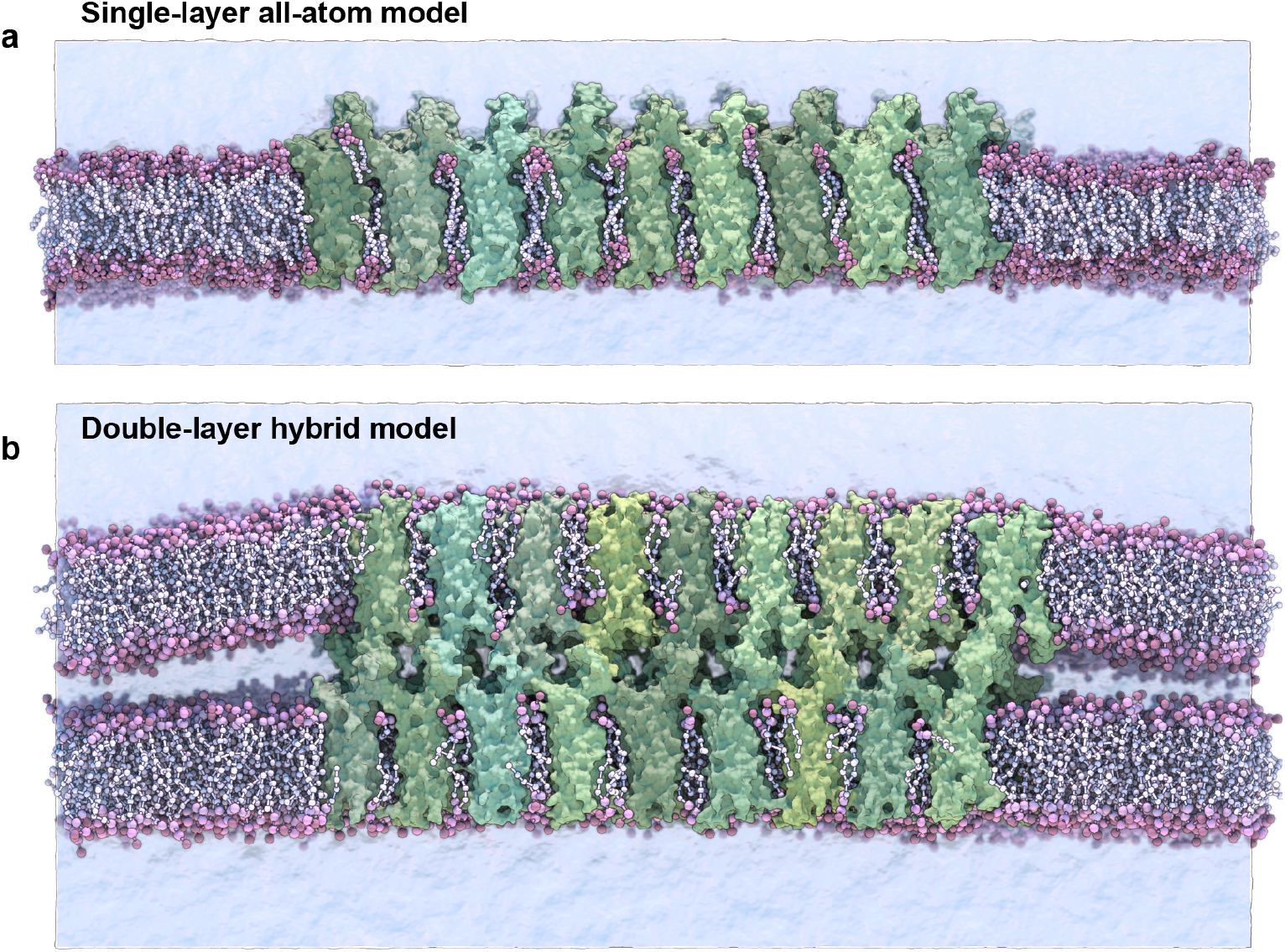
Single-layer and double-layer claudin-15 strands. (a) All-atom model of the single-layer claudin-15 strand in a single POPC lipid bilayer including 18 monomers (30 nm). (b) Double-layer claudin-15 strands embedded in two parallel POPC lipid bilayers represented at hybrid resolution with 36 monomers and 18 paracellular channels (30 nm). Additional systems of claudin-15 strands with 63 nm and 135 nm lengths were also simulated.

The initial structure of wild-type (WT) claudin-15 strands were based on our refined model of claudin-15 channels with twelve monomers embedded in two parallel lipid membranes^23^. The “single-layer” models of claudin-15 in the POPC bilayer were constructed by selecting a single layer of claudin strands with their surrounding lipid bilayer in the refined model and replicating it three times along its length to build a 30 nm claudin-15 strand with 18 monomers (Fig. 1a). Two patches of POPC lipid bilayers (20 nm×10 nm) were added on the two ends of the system to separate the system from its periodic images. The system was then solvated and neutralized with 150 mM of NaCl salt via the molecular visualization program VMD^39^. The resulting system consists of 18 claudin-15 monomers, 2045 lipid molecules, 183,888 waters, and 1038 Na^+^ and Cl^−^ ions in a 20 nm×50 nm×26 nm simulation box. The A134P mutant system was prepared by mutating the alanine (A) at position 134 on the claudin-15 sequence to proline (P) in this model using VMD^39^. The WT and the A134P mutant single-layer systems were simulated at all-atom resolution for 547 ns as described below. In the first step, the protein backbone and lipid head groups were restrained harmonically with a force constant of 2 kcal/mol.Å^2^ and the lipid tails were equilibrated for 200 ps at constant volume and temperature. In the next step, the lipid head groups were released and the system was equilibrated at constant pressure for 2 ns, after which, the two extracellular loops of claudins (residues 33–44 of ECS1 and residues 148–155 of ECS2) were released to move freely while the rest of the protein backbone was gradually released by decreasing the force constant to 1.0, 0.75, 0.5, and 0.25 kcal/mol.Å^2^ over 45 ns. After releasing all restraints, the system was equilibrated freely for 500 ns at constant pressure.

Two sets of claudin-15 strands corresponding to the WT and A13P mutant were prepared in a double-membrane system (here referred to as the “double-layer” systems). The initial structure of the double-layer systems was the refined model of claudin-15 channels with twelve monomers (3 claudin-15 channels)^23^, which was replicated along the strand to create 30 nm, 63 nm, and 135 nm-long strands. These systems consist of 36, 84, and 180 claudin monomers in two parallel POPC lipid membranes (Fig. 1b). The ions in the equilibrated refined model of the three-channels system (150 mM NaCl) were replicated along with the protein and lipid bilayers to construct the three large systems. Four patches of 20 nm×10 nm POPC lipid bilayers were added at the two ends of the system to separate the claudin strands from their periodic images. The corresponding A134P mutant systems for each strand were then generated in VMD^39^ by mutating the alanine (A) at position 134 to proline (P).

The six double-layer systems were then converted into hybrid-resolution models using the PACE forcefield^40–42^. PACE forcefield models lipids and solvents consistent with the MARTINI forcefield^43^, while proteins are represented by a united-atom (UA) model, where heavy atoms and polar hydrogens are explicitly represented. The conversion was performed using the python scripts provided by Han’s laboratory at (http://web.pkusz.edu.cn/han/) to convert protein, lipids, and ions into the hybrid model. The resulting system was then solvated by adding MARTINI water molecules^43^. The hybrid model systems ranging from 20 nm×50 nm× 26 nm to 20 nm×150 nm× 26 nm consist of 200K-800K particles.

The six double-layer systems were equilibrated for 1.125 *µ*s following a process similar to the equilibration of the all-atom system. Briefly, by harmonically restraining the protein backbone and lipid head groups with a force constant of 2 kcal/mol.Å^2^, the lipid tails were equilibrated for 200 ps at constant volume and temperature. The lipid head groups were then released and the systems were equilibrated for 2 ns at constant pressure and temperature. Finally, the two extracellular loops of claudins (residues 33–44 of ECS1 and residues 148–155 of ECS2) were released to move freely while the rest of the protein backbone was gradually released by decreasing the force constant to 1.0, 0.75, 0.5, and 0.25 kcal/mol.Å^2^ over 70 ns. The equilibration process was performed with a time step of 2 fs. The simulations were then followed by 1.055 *µ*s of simulation time at constant pressure, using a time step of 2 fs for the first 140 ns and a time step of 3 fs for the last 915 ns.

### Molecular dynamics simulations

MD simulations were performed using the program NAMD^44^. All-atom simulations were carried out using CHARMM36 forcefield for proteins^45–47^, ions^48^, and phospholipids^49^ with the TIP3P water model^48^. These simulations were carried out with a time-step of 1 fs and assuming periodic boundary conditions. Langevin dynamics with a friction coefficient of *γ* = 5 ps^−1^ was used to keep the temperature constant at 333 K. The Langevin Nosé-Hoover method^50^ was used to maintain the pressure at 1 atm in constant pressure simulations. Long-range electrostatic forces were calculated using the particle mesh Ewald method^51^ with a grid spacing of at least 1 Å in each direction. The simulations used a time-step of 1 fs, 2 fs, and 4 fs for bonded, short-range nonbonded, and long-range electrostatic interactions calculations, respectively. A 1-4 rule is applied to the calculation of nonbonded interactions. Additional restraints were applied by enforcing a harmonic potential with a force constant of 2.0 kcal/mol.Å^2^, unless otherwise stated.

The hybrid resolution PACE models were simulated assuming periodic boundary conditions and using a time-step of 2 fs or 3 fs. The dielectric constant is set to 15 as recommended for MARTINI simulations with non-polarizable water^43^. The electrostatic interactions are switched to zero between 9 Å and 12 Å, and the van der Waals interactions cutoff is set to 12 Å. Langevin dynamics with a friction coefficient of *γ* = 5 ps^−1^ was used to keep the temperature constant at 333 K. The Langevin Nosé-Hoover method^50^ was used to maintain the pressure at 1 atm in constant pressure simulations.

### TM3 tilt and bending angle calculations

The tilt angle of the third transmembrane helix (TM3) in claudins is defined as the angle between the helical axis and the bilayer normal. The helical axis of TM3 in all-atom simulations of WT and A134P mutant was calculated using the HELANAL algorithm^52–55^ and was averaged over the last 100 ns of the trajectories with a frequency of 60 ps. The bending angle of TM3 at position 134 is defined as the angle between the two vectors connecting A134/P134 to the two ends of the TM3 helix. The two points defining the endpoints of TM3 are residue 123 on the cytoplasmic side and residue 141 on the extracellular side of TM3. The position of C*α* of these residues is used to calculate the bending angle, which is averaged over the last 100 ns of the trajectories with a frequency of 60 ps.

### Curvature calculations

The local curvature of the strands in the plane of the membrane is calculated for the longest double-layer system (135 nm with 180 claudins) for the WT and the A134P mutant. The local curvature is estimated as the reciprocal of the radius of tangential circles fitted to the strands. For these calculations, claudin strands are projected into a two-dimensional polymer in the plane of the membrane. The center of mass of the each four pore-forming claudins constitutes a data point *i* marked as (*x*_*i*_, *y*_*i*_). For the two 135 nm-strands, 44 data points are recorded. To determine the radius at each position *i*, a circle is fitted to the data points between *i*-5, *i*+5 (for *i* = 6, …, 39). The best fit is determined via least-square linear regression and the local radius *r*_*i*_ is used to calculate the curvature 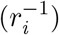. For each system, the last 600 ns of the simulation trajectories with a frequency of 60 ps are used for these calculations. Twelve data points from the ends of the strands were excluded from these calculations to exclude the effect of interaction of the strands with their images across the periodic boundaries.

### Persistence length calculations

We calculated the persistence length (*l*_*p*_) of claudin-15 strands from equilibrium MD simulation trajectories. We used the worm-like chain approximation^56–58^ to estimate the persistence length of the strands in the plane of membrane from thermal fluctuation in the strands. In these calculations, claudin strands are considered two-dimensional linear polymers in the plane of the membrane that are made of discrete segments. Each four pore-forming claudins are considered to be a segment located at position 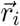, where 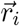 is the center of mass of the four claudins.

At each point on the strand 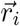, the tangent angle *φ*_*i*_ is determined as the angle corresponding to the tangent vector 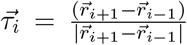. The persistence length is defined as the distance along the strand over which the tangent angles become uncorrelated. Assuming a discrete strand made of segments with length *δs*, the contour length between any two points *i* and *j* on the strand is estimated to be *s* = |*i* − *j*|*δs*. We defined the contour fragment *δs* to be the distance between two segments 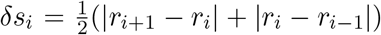 averaged over all segments during the simulation time; 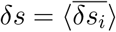.

The persistence length *l*_*p*_ is then calculated as the length over which the correlation between the tangent angles defined as (cos (*φ*(*s*) − *φ*(0))) is dropped by a factor of e:

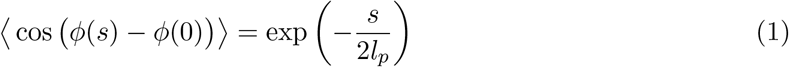

The factor of two in the denominator is due to the two-dimensional nature of the strands. We calculated the persistence length for three WT strands and the three A134P strands in the double-layer systems with lengths ranging from 30 nm to 135 nm. The last 600 ns of the simulations trajectories were used for this calculation with a frequency of 60 ps. The slope of the initial decay of the logarithm of the cosine correlation functions is used to calculate the persistence length. The number of points included in curve fitting is determined by an R-squared cut-off of 0.95.

## Results and discussion

In order to investigate the impact of TM3 structure on the flexibility of claudin-15 strands, we carried out MD simulations of the wild-type (WT) and A134P mutant claudin-15 strands in double- and single-layer membrane systems (Fig.1). These models of claudin strands were generated based on our previously refined structure of claudin-15 strands^23,38^ in accordance with the proposed architecture of Suzuki et al^24^. Eight systems of various lengths, ranging from 30 nm to 135 nm with 26 to 180 claudin monomers were studied. Due to the large size of the systems and the time scales required to capture the strand fluctuations (~ *µ*seconds), we carried out the double-membrane strand simulations at a hybrid resolution using the PACE forcefield^40–42^. The PACE forcefield combines a united atom (UA) representation of proteins with a coarse-grained (CG) model of lipids and solvents to efficiently capture the dynamics of the strands at large scales^38,59–62^, allowing us to calculate the persistence length of the WT and mutant strands and compare their lateral flexibility. To assess the effect of A134P mutation on conformational changes of the protein and side-by-side *cis-*interactions between claudin monomers, the WT and A134P mutant strands were simulated at atomic resolution in a single lipid membrane. These simulations enabled us to compare the secondary structure of TM3 in WT and A134P mutant strands within a more realistic environment and to perform a more detailed analysis of the side-by-side (*cis-*) interactions between neighboring monomers.

### Claudin-15 strand stability

We assessed the structural stability of claudin-15 strands in “single-layer” and “double-layer” membrane systems simulated at various lengths. The two single-layer systems of WT and A134P mutant were stable during the 500 ns simulations. The average root mean square deviation (RMSD) of claudin-15 backbones with respect to their initial structures was 3.0±1.0 Å for the WT strand and 2.8±0.5 Å for the mutant strand. Interestingly, claudin monomers were slightly more stable in the mutant strand. The RMSD of claudin pairs with respect to their initial conformation, an indication of the stability of the-side-by-side interactions, was also comparable for the WT and mutant strands (3.2±0.6 Å for the WT and 3.0±0.3 Å for the mutant backbones).

The double-layer systems simulated at hybrid resolution were stable during the 1*µ*s simulation trajectories with an average RMSD of 4.3±0.1 Å for the WT and 4.4±0.1 Å for the mutant strands with respect to the initial structure. The protein stability is comparable in all-atom and the hybrid-resolution models. We have previously reported a detailed comparison of structural properties of claudin strands in hybrid-resolution and atomic simulations showing similar stabilities of two models^38^. In the double membrane systems, the tetrameric claudin channels formed between the two membranes were also stable with an average RMSD of 6.7±0.3 Å for the WT and 6.6±0.2 Å for the mutant strand backbone with respect to the initial structures.

### A134P mutation reduces lateral flexibility of claudin-15 strands

The A134P mutant is expected to affect the two-dimensional rigidity of claudin strands in the membrane^21^. We calculated the persistence length of claudin strands in the double-layer systems and compared it between the WT and mutant strands. The persistence length (*l*_*p*_) is an indication of lateral flexibility of the strands in the plane of membrane and is directly related to their bending rigidity as well as the temperature^38^. The persistence length was calculated for the six double-layer systems of WT and A134P mutant with lengths ranging from 30 nm to 135 nm. The calculations are similar to our previous work^38^, and are briefly described in the Methods section. In summary, *l*_*p*_ is calculated as the distance over which the direction of the strand becomes uncorrelated and is estimated from exponential decays of the correlation function of tangent vectors over the length of the strand. We calculated the persistence length from thermal fluctuations of the strand in equilibrium simulations over *µ*-long trajectories (Fig. 2a,b). The persistence length of the WT strands was 187.8 nm with a standard deviation of 38.9 nm across the three systems, while the A134P mutant strands had an average persistence length of 732.7 nm with a standard deviation of 126.8 nm. The WT persistence length is consistent with our previously reported values for claudin-15 (~174 nm)^38^, and falls within experimentally estimated values of claduin-15^35^. The only experimental measurement of the persistence length in claudin-15 is from analyses of local curvature of the strands in freeze-fracture electron microscopy of the WT claudin-15 by Zhao et al^35^ with a relatively large error bar (191 nm±184 nm).

**Figure 2:**
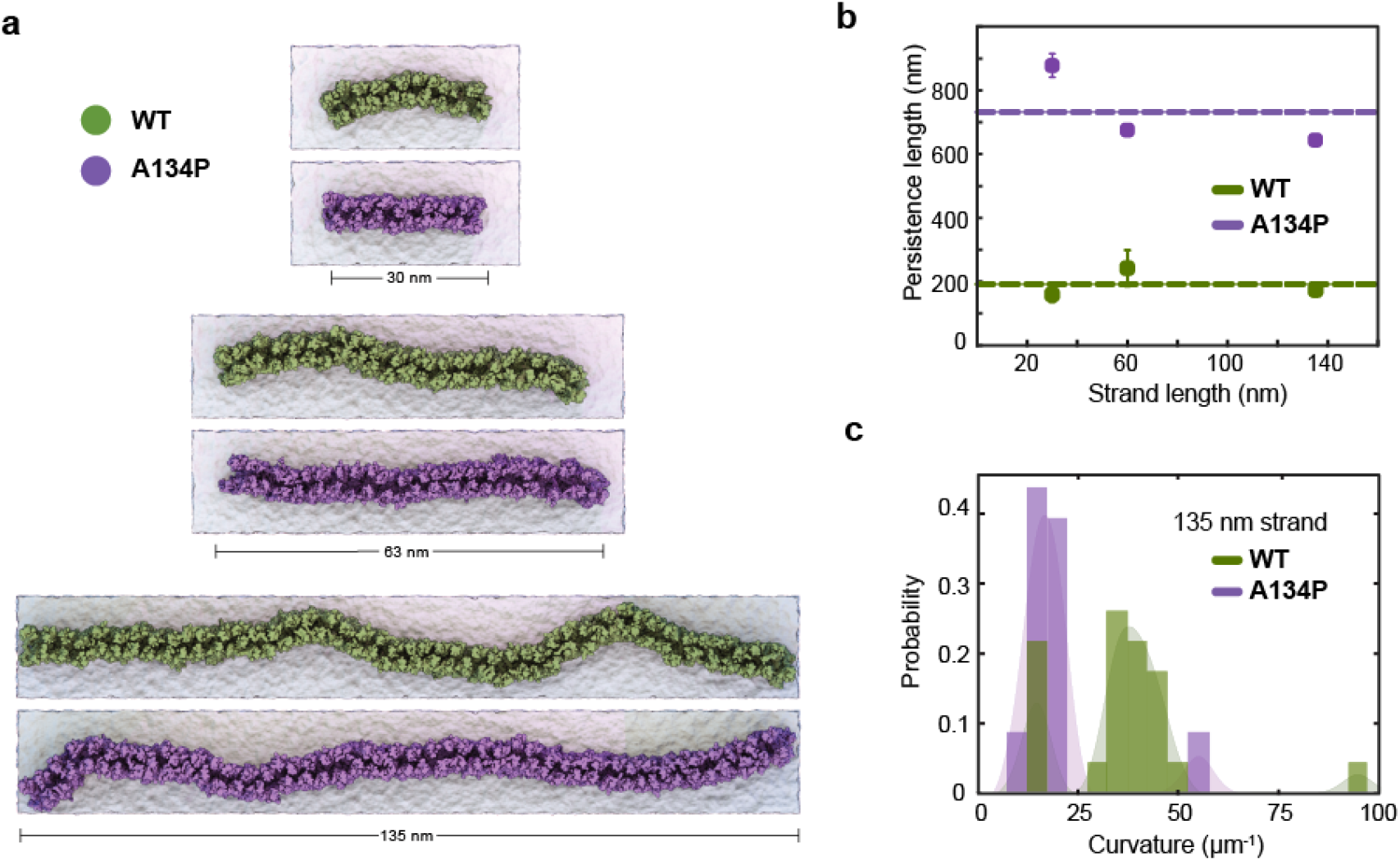
TM3 bending restrict claudin-15 strand flexibility. (a) Equilibrated configuration of WT and A134P strands of various lengths simulated for 1 *µ*s in double parallel lipid bilayers and hybrid resolution, show the “curvy” morphology of cluadin-15 strands as opposed to the more “straight” shapes of A134P mutant strands. (b) The strand persistence length is calcualated for each length of WT and A134P claudin-15 strands. The horizontal lines show the average persistence length for WT and A134P mutant strands. (c) The distribution of local curvature along the length of the longest simulated strand (135 nm) is calculated for WT and A134P strands.

The larger persistence length of the A134P mutant strands in comparison to the WT claudin-15 strands (~3.5 fold) indicates that the A134P mutants are less flexible with a relatively high bending modulus. The bending rigidity of the strands, and thus, their persistence length is directly related to the average curvature of the strands over time^38^. We calculated the average curvature of the strands over time for the longest (135 nm-long) strand simulated here for the WT and the A134P mutant (Fig. 2c). The average local curvature of the WT strand is approximately 36.5 *µ*m^−1^, while the local curvature of the A134P strand is significantly lower with a sharper distribution averaged at 19.8 *µ*m^−1^. The ~3.5 fold increase of the persistence length in the A134P mutant in our simulations is consistent with the lower curvature of the mutant strands in comparison to the WT strands (Fig. 2b,c) and indicates that the mutation significantly reduces the lateral flexibility of claudin-15 strands.

The A134P mutant is similar structurally to claudin-3, which is shown to have a distinct architecture with straight strands^21^. Interestingly, mutation of P134 in claudin-3 to an alanine (A), as in claudin-15, reproduces the signature curvature of claudin-15 strand morphology^21^. However, this site of interest had not yet been investigated in claudin-15. These simulations indicate that claudin-15 significantly decreases bending flexibility of the strands similar to those observed in freeze-fracture electron microscopy of claudin-3^21^, suggesting that residue 134 plays a key role in lateral flexibility of the strands. However, we can not yet rule out the role of other transmembrane residues in claudin assembly^26^ and their contribution to strand flexibility.

As we showed in our previous work, claudin-15 strand flexibility is mostly attributed to flexible *cis-*interfaces between neighboring claudins^38^. In particular, we showed that there are three sets of side-by-side *cis-*interaction between claudins pivoted at the extracellular helix (ECH), which confer flexibility to the strands. Residues 143-147 located on the extracellular end of TM3 play a key role in defining one set of these interfacial interactions. Mutation of P134 of TM3 in claudin-3 is shown to affect the secondary structure of TM3, and thus, it is not surprising that it affects the strand flexibility. To understand the molecular mechanism of strand flexibility and the effect of A134 mutation on the interfacial interactions, we investigated the secondary structure of TM3 in the all-atom simulations of WT and A13P mutant claudin-15 strands and analyzed the frequency of side-by-side *cis-*interactions and its implications on strand flexibility.

### A134P mutation results in TM3 bending

TM3 is the longest transmembrane helix in claudin-15 with 36 residues and extends beyond the membrane into extracellular solution (Fig. 3a). In claudin-15, A134 is located in the middle of TM3 near the extracellular lipid-water interface. A proline at the position of this residue is expected to create a bend in the TM3 helix as observed in the crystal structure of claudin-3^21^ and claudin-4^20^ and transmembrane helices of other membrane proteins^63,64^. This is while the structures of claudin-15^18^ and claudin-19^19^ with alanine at this position, and P134A/G claudin-3 mutants^21^ show a relatively straight conformation of the TM3 helix.

**Figure 3:**
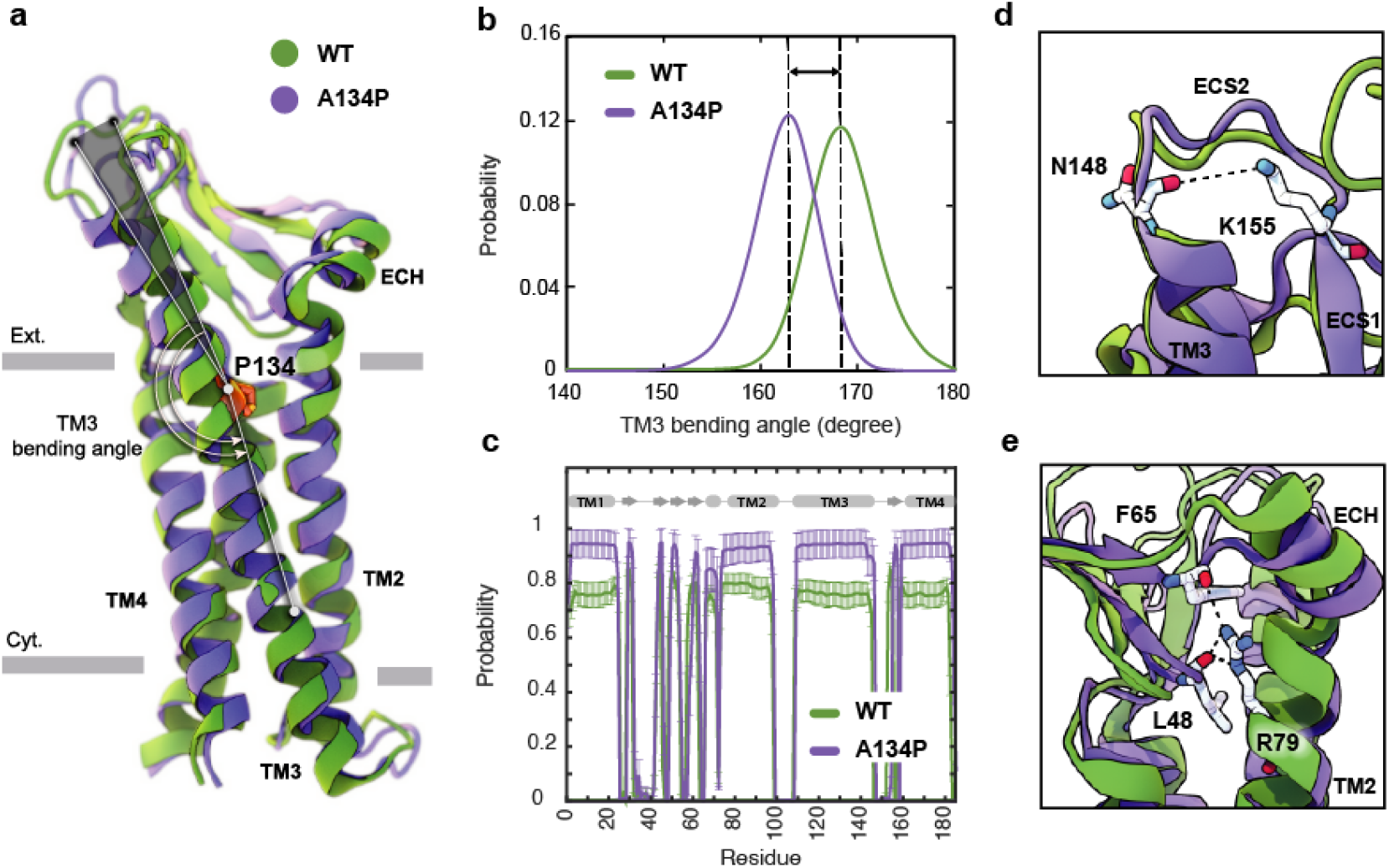
A134P mutation results in the bending of TM3. (a) Superposition of WT and A134P mutant claudin-15 monomers exhibits the bent structure of TM3 after A134P mutation. Gray bars suggest boundaries of the outer (Ext.) and inner (Cyt.) leaflets of the lipid bilayer. (b) TM3 bending angle for the WT and A134P mutant monomers is calculated over the last 100 ns of the simulation trajectories of single-layer strands. (c) The secondary structure of helices and *β*-strands in WT and A134P monomers show structural stability of the monomers over the last 250 ns of simulation time. Key interactions within each monomer, in the form of (d) a salt bridge between N148 and K155 stabilizes the structure of ECS2, and (e) interactions between the backbone of F65 and L48 *β*-strands and side chain of R79 on TM2 stabilize the *β*-sheets and orientation of ECH. These interactions are maintained in mutant monomers (shown in purple) similar to WT monomers (shown in green).

Simulation trajectories of claudin-15 strands in a single membrane indicate that, in the WT strand, TM3 is helical with a helical axis that is titled by 8.3^°^ ± 2.2^°^ with respect to the membrane normal. Upon mutation of A134P, while the tilt angle of the helix does not change in the membrane, the mutation results in bending of the helix at position 134, where the extracellular end of the helix bends toward the membrane. The average bending angle of TM3 changes from 168.4^°^ ± 3.5^°^ in WT claudin-15 to 162.5^°^ ± 3.5^°^ in A134P mutant. The shift in the bending angle of claudin-15 after A134P mutation (~ 6^°^ difference) is comparable to the the difference observed between WT and P134A/G mutants claudin-3 (~ 8^°^ difference)^21^.

Superposition of the WT and A134P monomers shows that in the A134P mutant, the extracellular domains including, ECS1 with five *β*-strands, the loop between *β*1 and *β*2, and the extracellular helix (ECH), and ECS2 are inclined toward the membrane in the direction of the TM3 bending (Fig. 3a). Simulation and analyses of the extracellular domain of claudins indicate that these domains are relatively rigid and form the backbone of paracellular pores^22,23^. In claudin-15, the extracellular domains are directly connected to TM3 by residues 150-154 of the ECS2 loop. The conformation of ECS2 is stabilized by salt bridge between K155 and N148 (Fig. 3d), as identified in the crystal structure and MD simulations of claudin-15^18,23^ and homology models of claudin-5^28^. This interaction was conserved in ~ 83% of the simulation time in A134P mutant strand monomers and in ~ 86% of the time in WT strand monomers.

Another key interaction for stabilizing the structure of extracellular domain in claudin-15 is between the sidechain of an extended arginine R79 on TM2 and the backbone of F65 on ECH and L58 on the *β*-sheets (Fig. 3e). This interaction which was observed in the crystal structure of claudin-15^18^ is important in stabilizing the extracellular *β*-sheets and keeping the short extracellular helix ECH in place, i.e. parallel to the membrane and perpendicular to TM2. This set of interactions was maintained for 97% of the simulation time in A134P mutant monomers and 85.5% in WT monomers. It is through these robust interactions that the extracellular domains maintain a rigid conformation that is critical for the tetrameric assembly of claudins into paracellular pores.

Despite the bent structure of TM3 and the shift in extracellular domains positioning, the overall secondary structure of mutant monomers was maintained 91.6% of the simulation time for helices and 75.9% for *β*-strands in A134P mutant monomers, compared with 83.8% and 78.3% for the WT monomers (Fig. 3c). As indicated by these results, mutant monomers are more stable and rigid in their structures compared with WT monomers. The rigidity of the structure of mutant monomers is projected into mutant strands dynamics, with lower fluctuations in their shapes and minimal deviations of monomers from the straight arrangement. To explain the effect of bent TM3 and the rigidity in mutant monomers on strands flexibility further, we explored the flexibility of *cis-* interactions in WT and mutant monomers.

### Effect of TM3 bending on *cis-*interactions

The linear arrangement of claudins in a row in the membrane is maintained through two sets of *cis-*interactions; a set of side-by-side *cis-*interactions as observed in the crystal structure of claudin-15^18^, as well as a set of “face-to-face” *cis-*interactions between two anti-parallel claudin rows originally proposed by Suzuki et al.^24^. TM3 is one of the main contact points between neighboring claudins. Residues 146 and 147 on the extracellular end of TM3 are critical for side-by-side *cis-*interactions and strand formation^18,23,28,65^.

We have recently proposed a model for claudin strand flexibility, in which the lateral flexibility of the strands is attributed to flexible side-by-side interactions between claudin monomers^38^. In this model, claudin-15 monomers move as tetrameric units within the strands. In this movement tetrameric units forming ion channels slide with respect to each other to adjust to the local curvature of strands imposed by external restraints. Those simulations revealed three dominant side-by-side interfaces pivoted at the extracellular helix ECH (residues 66-71) as claudins monomers rotate/slide with respect to each other within the strand. Interactions between ECH (S68) and the extracellular end of TM3 (F146) (Fig. 4a) are one of the three interfacial interactions and are dominant at zero or positive (outward) curvatures of the strands. The second set of interactions is between ECH (S67) and ECS2 (E157) (Fig. 4a) that are again mostly present at zero or positive curvatures of the strands^38^. At negative (inward) curvatures, both of these interaction sets are weakened and are replaced by interactions between ECH and ECS1.

**Figure 4:**
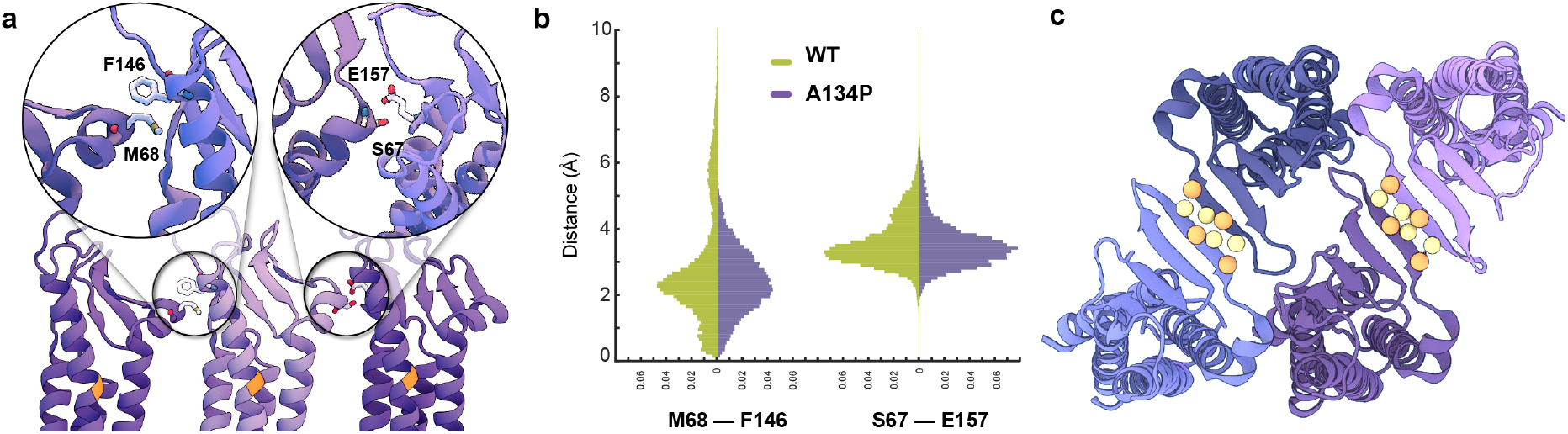
Side-by-side *cis*-interactions between WT and A134P claudin-15 strands. (a) A snapshot of three claudin-15 monomers in the single-layer strands after 500 ns of simulation in all-atom resolution highlighting two sets of side-by-side *cis*-interactions. (b) Pair-wise distance distribution of M68 (ECH)–F146 (TM3) and S67 (ECH)–F157 (ECS2) in the all-atom simulation of the single-layer WT and A134P mutant claudin-15 strands. The extended tail of the distribution in the case of WT suggests more flexible side-by-side interactions in the WT compared to the A134P mutant. (c) “Face-to-face” interactions between claudins are maintained through hydrogen bonds between the backbone of the *β*-sheets of claudins. These interactions are maintained throughout the simulation of both WT and mutant strands. The position of *C*_*α*_ of residues involved in hydrogen-bonding is highlighted as spheres.

Analyses of our all-atom simulation trajectories indicate that both ECH-TM3 and ECH-ESC2 interacting interfaces are present in the WT and A134P claudin-15 strands (Fig. 4b). However, the distribution of the pair-wise distances between the residues involved in these two sets differs between the WT and A134P mutant. The extended distribution of pair-wise distances (up to 10Å) in the WT claudin-15 strand indicates loose interactions between neighboring monomers and a more flexible *cis-*interface. The extended tail of the distribution corresponds to instances in which these interactions are replaced with those involving ECH and ECS1 dominant at negative (inward) curvatures. In contrast, the sharp distance distribution of pairwise distances (up to 6Å) in the A134P mutant indicates tight interactions and therefore a more rigid configuration for the strands. In the WT trajectories, the TM3-ECH interface represented here by M68-F146 interaction is only present in ~82 % of the simulation time, while the same interaction is maintained for ~95 % of the simulation time in the A134P mutant. The TM3-ECH interactions are mostly present at zero (straight) or positive (outward) curvatures and are expected to vanish at negative (inward) curvatures^38^. Similarly, the interactions between ECH and ECS2 (S67 and E157) is less maintained in the WT compared to the mutant. Both of these interactions are dominant in the straight configuration of A134P mutant consistent with our previous results^38^, however, the configurations dominant at negatively curved strands corresponding to the extended tail of ECH-TM3 distance distribution are only present in the WT strand. This indicates that mutation of A134P in TM3 strengthens the interactions between TM3 and ECH, and limits its replacement with alternative interactions including ECS1. Thus, these simulations provide the molecular basis of the increased persistence length and decreased flexibility of claudin-15 strand upon A134P mutation.

While side-by-side *cis-* interactions demonstrated variations between the WT and A134P mutant strands, other inter-molecular interactions were well-maintained. The face-to-face interactions formed between the claudins in opposing rows of the same membrane were not affected by the A134P mutation. The antiparallel double-row arrangement of claudins in a single lipid membrane is stabilized through hydrogen bonds between two *β*4 strands of neighboring claudins (Fig. 4c). The average distance between the *β*-strands of claudins over the last 15 ns of the simulation was comparable for the WT (3.34 ± 1.52 Å) and mutant (2.83 ± 0.03 Å) systems, indicating that face-to-face interactions were not affected by the TM3 bending in A134P mutant strands.

These simulations corroborate the results of the double-layer systems by indicating that the A134P mutation reduces the lateral flexibility of claudin-15 strands. The broader range of *cis*-interfaces observed in WT strands allowed interacting monomers in the same membrane to rotate relative to their neighboring monomer, resulting in higher curvature of the WT strands. Conversely, the *cis*-interfaces became rigid due to a bent TM3 in A134P mutant claudins, which explains the lower flexibility of the A134P mutant and the more straight shape of tight junction strands in claudin-3^21^. A tight *cis-*interface limits relative movement of claudins and inhibits bending of the strands (to negative curvatures).

## Conclusions

We investigated mechanical flexibility of claudin-15 strands and the effect of a single point mutation A134P on lateral flexibility of the strands. The A134 residue is located on the third transmembrane helix (TM3) of claudin-15 and is not directly involved in claudin-claudin interactions. However, it was recently suggested that the distinct morphology of tight junctions formed with claudin-3 is due to a proline at this position, and mutation of P134 to an alanine (A) in claudin-3 resulted in strands similar to claudin-15^21^. Here, we investigated the reverse effect in claudin-15, i.e. mutation of A134 to proline (P) in claudin-15 via MD simulations. Our results reveal that the A134P mutation increases the persistence length of claudin-15 by more than 3x, consistent with the comparatively straight shape of claudin-3 strands^21^.

Our results indicate that the A134P mutation does not establish any new contacts between neighboring claudins nor does it eliminate any of the previously reported contacts^18,38^. However, it does change the dynamics of the strand fluctuations. In a recent model describing the dynamics of claudin-15 strands, we showed that lateral flexibility of claudin strands is due to flexible side-by-side *cis-*interactions pivoted at the short extracellular helix ECH^38^. Mutation of A134 to proline locks this pivoted movement to switch between two out of three interfaces with limited occurrence of the third interface. Considering that the strands are made of an *anti-parallel* double row of claudins, this subtle change results in a relatively straight shape of the mutant strands. The mutation might also indirectly affect other potential *cis-*interfaces to influence strand morphology^26,33,66,67^.

These findings corroborate the role of *cis-*interactions in conferring flexibility to the strand and their effect on strand morphology. They elucidate the indirect role of transmembrane helices, not necessarily on claudin assembly, but on modulating their dynamical properties by providing a microscopic description for large-scale properties of the strand at micro-meter length scales.

## Acknowledgements

The work of F.K.-A., S.F., S.M. was supported by the National Science Foundation grant MCB-1846021. This research is part of the Frontera computing project at the Texas Advanced Computing Center. Frontera is made possible by National Science Foundation award OAC-1818253.

## Author contributions

All authors contributed to the writing of the manuscript. F.K.-A and S.F. designed the research.

S.F. carried out the simulations. S.F. and S. M. analyzed the data and prepared the manuscript.

